## Supplemental tables 1-3 for "Bacterial DNA on the skin surface overrepresents the viable skin microbiome"

**Supplemental information**

| **Name** | **Description** | **Sequence (5' - 3')** | **Reference** |
| --- | --- | --- | --- |
|  | ddPCR FP (Universal bacterial 16S qPCR FP) | TCCTACGGGAGGCAGCAGT | (*37*) |
|  | ddPCR RP (Universal bacterial 16S qPCR RP) | GGACTACCAGGGTATCTAATCCTGTT | (*37*) |
| EFTU_FP | Staph-specific ddPCR forward primer | ATGCCACAAACTCGTGAACA | this paper |
| EFTU_RP | Staph-specific ddPCR reverse primer | ACATCGTCACCTGGGAAGTC | this paper |
| EUB338 | Pan-bacterial FISH probe | GCTGCCTCCCGTAGGAGT | (*17*) |
| NonEUB338 | Nonsense control FISH probe | CGACGGAGGGCATCCTCA | (*38*) |
|  | C. acnes FISH probe | GAGTGTGTGAACCGATCATGTAGTAGGCAA | (*39*) |
| 27F | Forward 16S sequencing primer | AGAGTTTGATCCTGGCTCAG | (*40*) |
| 534R | Reverse 16S sequencing primer | ATTACCGCGGCTGCTGG | (*40*) |

**Table S1.** Nucleotide sequences used in this study.

| **Sample** | **Sequence count** |
| --- | --- |
| **HV4 Nares +PMA** | **153847** |
| **HV1 Nares +PMA** | **114649** |
| **HV1 RAC +PMA rep1** | **108308** |
| **HV2 Glabella +PMA** | **101272** |
| **Hair shaft +PMA rep2** | **97416** |
| **HV3 Glabella -PMA** | **76165** |
| **HV1 Glabella -PMA rep4** | **70341** |
| **HV2 Nares -PMA** | **69549** |
| **HV2 Glabella -PMA** | **67565** |
| **HV1 Glabella -PMA rep1** | **61693** |
| **HV1 TW -PMA** | **61628** |
| **HV3 Glabella -PMA** | **60889** |
| **HV3 RAC -PMA** | **60330** |
| **HV2 AF -PMA** | **59564** |
| **HV1 RAC -PMA rep2** | **59158** |
| **HV2 PF -PMA** | **58842** |
| **HV1 Glabella -PMA rep3** | **56854** |
| **HV4 RAC -PMA** | **54039** |
| **HV2 Back -PMA** | **53930** |
| **HV3 Nares +PMA** | **53143** |
| **HV2 VF -PMA** | **52215** |
| **Hair shaft -PMA** | **51698** |
| **HV3 RAC -PMA** | **50870** |
| **HV4 Glabella -PMA** | **49799** |
| **HV4 RAC +PMA** | **49501** |
| **HV2 RAC -PMA** | **49497** |
| **HV3 VF -PMA** | **48957** |
| **HV3 Nares -PMA** | **48546** |
| **HV3 Nares +PMA** | **48511** |
| **HV4 Nares -PMA** | **47917** |
| **HV3 Back -PMA** | **46631** |
| **HV1 Nares -PMA** | **46250** |
| **HV3 RAC +PMA** | **45487** |
| **HV3 AF -PMA** | **43929** |
| **HV2 Nares +PMA** | **43324** |
| **HV1 RAC -PMA rep1** | **42463** |
| **HV2 RAC +PMA** | **40333** |
| **HV1 Back -PMA** | **39487** |
| **HV3 nares -PMA** | **39419** |
| **HV1 Glabella +PMA rep4** | **39357** |
| **HV1 Glabella +PMA rep2** | **38231** |
| **HV1 TW +PMA** | **36139** |
| **HV1 Glabella -PMA rep2** | **35249** |
| **HV3 RAC +PMA** | **34519** |
| **HV1 Glabella +PMA rep3** | **34273** |
| **HV1 Back +PMA** | **34201** |
| **HV3 Glabella +PMA** | **32629** |
| **HV3 PF -PMA** | **31665** |
| **HV1 Glabella +PMA rep1** | **26953** |
| **HV1 RAC +PMA rep2** | **25388** |
| **HV1 PF -PMA** | **20970** |
| **Hair shaft -PMA rep2** | **18791** |
| **HV4 Glabella +PMA** | **18636** |
| **HV1 PF +PMA** | **17414** |
| **Hair shaft -PMA rep1** | **17389** |
| **HV3 Back -PMA** | **17115** |
| **HV3 AF -PMA** | **16969** |
| **HV4 VF -PMA** | **16956** |
| **HV4 PF -PMA** | **15653** |
| **HV1 VF -PMA** | **15134** |
| **HV3 AF +PMA** | **14639** |
| **Hair shaft -PMA rep1** | **14192** |
| **HV3 VF +PMA** | **9156** |
| **Hair shaft +PMA** | **9136** |
| **HV1 AF -PMA** | **9096** |
| **HV4 PF +PMA** | **7483** |
| **HV3 PF +PMA** | **7438** |
| **HV2 Back +PMA** | **6296** |
| **HV4 AF -PMA** | **6261** |
| **HV3 PF -PMA** | **6154** |
| **HV2 VF +PMA** | **6032** |
| **Hair shaft +PMA rep1** | **5321** |
| **HV3 VF -PMA** | **5281** |
| **HV2 PF +PMA** | **5095** |
| **HV4 Back -PMA** | **4793** |
| **blank rep1** | **4781** |
| **HV3 Glabella +PMA** | **4593** |
| **Hair shaft +PMA rep1** | **4428** |
| **HV3 Back +PMA** | **4286** |
| **HV3 Back +PMA** | **4191** |
| **HV3 VF +PMA** | **4180** |
| **HV1 VF +PMA** | **3596** |
| **HV4 Back +PMA** | **3315** |
| **blank rep2** | **3291** |
| **Hair shaft -PMA rep2** | **3053** |
| **HV3 AF +PMA** | **2647** |
| **Hair shaft +PMA rep2** | **1819** |
| **HV4 VF +PMA** | **1408** |
| **HV3 PF +PMA** | **1111** |
| **HV2 AF +PMA** | **1033** |
| **HV4 AF +PMA** | **572** |
| **HV1 AF +PMA** | **421** |

**Table S2.** Sequence counts for Figures 2-4.

| **C. acnes** | **527328** |
| --- | --- |
| **Skin cocktail** | **500040** |
| **V2.3 -PMA** | **492270** |
| **V1.2 -PMA** | **455368** |
| **V3.0 -PMA** | **450136** |
| **V3.0 +PMA** | **446767** |
| **V1.1 -PMA** | **437976** |
| **V3.1 -PMA** | **434868** |
| **V4.3 -PMA** | **412972** |
| **V2.0 -PMA** | **403077** |
| **V3.2 -PMA** | **379443** |
| **V5.0 -PMA** | **376615** |
| **C. striatum** | **375535** |
| **V3.3 -PMA** | **366751** |
| **V2.2 -PMA** | **363327** |
| **V5.3 -PMA** | **338217** |
| **V2.1 -PMA** | **334685** |
| **S. epidermidis** | **328603** |
| **V1.1 +PMA** | **315313** |
| **V5.2 -PMA** | **313142** |
| **V1.2 +PMA** | **312954** |
| **V1.3 -PMA** | **311212** |
| **V1.3 +PMA** | **309985** |
| **V4.2 -PMA** | **307476** |
| **V4.0 -PMA** | **306340** |
| **V2.1 +PMA** | **298029** |
| **V1.0 +PMA** | **288799** |
| **V2.3 +PMA** | **285720** |
| **V4.2 +PMA** | **280293** |
| **V4.1 -PMA** | **255623** |
| **V3.1 +PMA** | **254358** |
| **V4.3 +PMA** | **248363** |
| **V1.0 -PMA** | **244772** |
| **M. luteus** | **233452** |
| **PBS 0 +PMA** | **227814** |
| **V5.0 +PMA** | **218913** |
| **V5.1 -PMA** | **217713** |
| **V2.0 +PMA** | **211122** |
| **V4.1 +PMA** | **200852** |
| **V5.2 +PMA** | **198699** |
| **V4.0 +PMA** | **197961** |
| **V3.2 +PMA** | **197419** |
| **PBS 3 -PMA** | **183592** |
| **V3.3 +PMA** | **175750** |
| **V2.2 +PMA** | **157845** |
| **PBS 2 -PMA** | **133797** |
| **PBS 0 -PMA** | **117178** |
| **PBS 3 +PMA** | **106895** |
| **PBS 1 +PMA** | **91483** |
| **V5.3 +PMA** | **51933** |
| **PBS 2 +PMA** | **35707** |
| **V5.1 +PMA** | **32808** |
| **PBS 1 -PMA** | **25336** |

**Table S3**. Sequence counts in perturbation recovery
